# REPRODUCTIVE SENESCENCE IMPAIRS THE ENERGY METABOLISM OF HUMAN GRANULOSA CELLS

**DOI:** 10.1101/2021.03.11.434795

**Authors:** Gustavo Nardini Cecchino, Juan A. García-Velasco, Eduardo Rial

**Author notes:** **Correspondence:** Department of Structural and Chemical Biology, Centro de Investigaciones Biológicas Margarita Salas, CSIC, Ramiro de Maeztu 9, 28040, Madrid, Spain. **Summary sentence:** Granulosa cells of women of advanced reproductive age show a reduced energy metabolism which leads to lower ATP levels.

## Abstract

Female age is the single greatest factor influencing reproductive performance. It is widely known that mitochondrial dysfunction plays a key role in reproductive senescence. Ovarian bioenergetics includes a sophisticated metabolic synergism between oocytes and human mural granulosa cells (GCs), which is crucial for oocyte maturation during follicular growth. These cells are believed to be potential biomarkers of oocyte quality. It has been proposed that alterations in their energy metabolism could lead to infertility. We investigated if there is an age-related effect on the energy metabolism of human mural granulosa cells. We performed an observational prospective cohort and experimental study including 127 women that underwent in vitro fertilization cycles allocated to two groups: a control group comprising oocyte donors aged less than 35 years and a group of infertile women aged over 38 years. The bioenergetics of cumulus cells and purified mural GCs were determined from oxidative phosphorylation parameters, aerobic glycolysis and adenine nucleotide levels. We have found that human mural GCs and cumulus cells present a high glycolytic profile and that the follicular fluid is critical to sustain their energy metabolism. GCs from older women present lower mitochondrial respiration and glycolysis than those from young donors which is not accompanied by a lower respiratory capacity. The diminished energy metabolism leads to a decrease in the total cellular energy charge. We conclude that, as women age, mural granulosa cells exhibit a reduction in their energy metabolism that is likely to influence female reproductive potential.

## Introduction

Female age is the single greatest factor influencing the reproductive performance of couples undergoing infertility treatments [1]. The 2016 final report from the Society of Assisted Reproductive Technology (SART) shows that nearly half of patients under 35 years undergoing an in vitro fertilization (IVF) cycle will achieve a live birth using their own eggs, whereas less than 26% of women aged over 37 years will. This becomes even more dramatic beyond 42 years-old with cumulative live birth rates as low as 3.7% [2]. Poor oocyte competence is the primary cause of age-related deterioration of reproductive capacity [3]. Indeed, oocyte donation cycles sustain a live birth rate of 50% irrespective of the women’s age [2].

Research on mitochondrial biology and cellular bioenergetics has gained increasing attention in recent years, providing new opportunities in different fields of medicine [4]. Thus, the role of mitochondria in cellular senescence and human aging has raised considerable interest [5–7]. It is known that mitochondrial dysfunction underlies some of the fertility defects in humans and that mitochondria play a critical role in oocyte quality and early embryo development [8–11]. In fact, mitochondria have been suggested to be a promising biomarker for IVF outcomes [12–13]. Furthermore, the possibility of overcoming mitochondrial dysfunction to improve oocyte quality and age-related infertility has become the goal of numerous studies [14–18].

Ovarian bioenergetics includes a sophisticated metabolic synergism between oocytes and granulosa cells (GCs), which is crucial for oocyte maturation during follicular growth [19,20]. However, little is known about human GC energy metabolism and its impact on IVF outcomes. In the field of assisted reproductive technology, previous studies that intended to explain defects in meiosis and other cellular processes affecting oocyte maturation, fertilization or embryo development have often focused either on the analysis of mtDNA as an indicator of mitochondrial content/function [8,10,21,22] or on ATP levels as an indicator of the cellular energy status [23–28]. Bioenergetics provides tools to analyse the factors that may lead to a decrease in the cellular energy levels. Therefore, we aimed to characterize the bioenergetic profile of human GCs in order to detect the potential impact of aging on the energy metabolism. We demonstrate that aged cells show a significant decrease of the two main ATP supply pathways leading to deficient cellular energy charge.

## Materials and methods

### Study design and population

We performed an observational prospective cohort including 127 women that underwent IVF and intracytoplasmic sperm injection after ovarian stimulation (OS) between December 2017 and December 2018 (Table 1). Patients were allocated to two age groups: the control group consisted of 84 oocyte donors aged less than 35 years, whereas the advanced reproductive age (ARA) group included 43 infertile women aged over 38. For the two groups, the following exclusion criteria were adopted: relevant systemic diseases, alcohol or drug abuse, heritable or chromosomal disorders, known mitochondrial dysfunction, anatomical defects of the reproductive system, formal contraindication to pregnancy or OS, prior exposure to radiation or chemotherapy, and a body mass index (BMI) ≥ 30 kg/m^2^. Among the diseases that could potentially impair fertility and mitochondrial function, we excluded in both groups: endocrine abnormalities, recurrent miscarriages, neurodegenerative and heart diseases, polycystic ovary syndrome, endometriosis, infectious and sexually transmitted disorders, and neoplasms. Thus, the group of women aged over 38 basically comprised infertile women due to male factor or unexplained infertility in which the presumable cause was the advanced age.

**Table 1.** Baseline characteristics, cycle parameters and ovarian response to OS

|  | **Control group** | **ARA group** | ***p*-value** |
| --- | --- | --- | --- |
| **Age (years)** | 23 (18-34) | 40 (38-46) | < 0.001 |
| **BMI (kg/m<sup>2</sup>)** | 22.4 ± 3 | 22.9 ± 2.7 | 0.323 |
| **AMH (ng/mL)** | - | 1.7 ± 1.7 | - |
| **AFC** | 19 ± 4.4 | 9 ± 4.2 | < 0.001 |
| **Estradiol levels (pg/mL)** | 2452 (288-9847) | 1729 (285-5844) | 0.08 |
| **Days of stimulation** | 11 (8-17) | 10 (7-16) | 0.711 |
| **Gonadotropin doses (IU)** | 2000 (775-4800) | 2250 (1200-4275) | < 0.001 |
| **Number of oocytes** | 19 (9-54) | 8 (2-25) | < 0.001 |
| **Number of mature oocytes** | 14 (5-41) | 6 (1-23) | < 0.001 |
| **Oocyte maturity rate** | 0.78 ± 0.13 | 0.80 ± 0.18 | 0.26 |

### Ovarian stimulation protocol and ovum pickup

Antagonist protocol was used to carry out OS in all cases. Individualized doses of gonadotropins were determined by an experienced specialist physician. Transvaginal ultrasound for cycle-monitoring was performed every 2 days starting on stimulation day 5. Fixed daily doses of 0.25 mg of gonadotropin-releasing hormone (GnRH) antagonist (Orgalutran, Organon; Cetrotide, Merck Serono; or Fyremadel, Ferring) were started when the leading follicle reached a mean diameter of 13-14 mm. All patients received GnRH agonist 0.2 mg (Decapeptyl, Ipsen Pharma) to achieve final oocyte maturation when the mean size of at least two follicles was 18 mm. Oocyte retrieval was performed 36 hours later under ultrasound guidance.

### Sample collection

Following oocyte retrieval, cumulus cells (CCs) were mechanically stripped from each oocyte using fine needles. Mural luteinized GCs were obtained from pooled follicular aspirate of follicles with at least 14 mm in mean diameter, as previously described [29] with minor modifications. Briefly, follicular fluid (FF) was slowly layered on 3:1 Ficoll gradient (Histopaque 1077, Sigma-Aldrich) and centrifuged at 400 g for 20 min. Follicular-derived cells were collected from the middle layer and washed twice with PBS. Cellular aggregates from both cell lines were dissociated using 0.5 mL of a 0.05% Trypsin-EDTA solution at 37°C for 5 min. 4.5 mL of our standard culture medium (M-199 supplemented with 10% heat-inactivated fetal bovine serum (FBS) and 100 U/mL of penicillin/streptomycin) was added to stop trypsin digestion, and the cell suspension was centrifuged at 500 g for 5 min. Subsequently, cells were incubated for 2 min at room temperature in the presence of red blood cell lysis buffer (Miltenyi Biotec S.I.) according to the manufacturer’s instructions. Finally, CCs and purified GCs were washed with PBS and resuspended in standard culture media. All the centrifugations were performed at room temperature. Occasionally, in order to reach the minimum cell number needed to perform the experiments, patient samples were pooled. However, it was always GCs within the same group of patients (control with control; ARA with ARA). The same procedure was followed, when necessary, with CCs. When collecting and preparing samples, aliquots of pooled follicular aspirates from each group of patients were centrifuged at 800 g for 20 min to remove cells. The supernatant consisted of purified FF which was sterile-filtered using a 0.22 μm pore size membrane filter (Minisart, Sartorius Stedim Biotech S.A.), alicuoted and stored at −80°C. A scheme with the complete experimental protocol is shown in Supplemental Figure 1.

### Assessment of the bioenergetic properties

Characterization of the bioenergetic properties of mural GCs and CCs was performed using an XF24 Extracellular Flux Analyzer (Agilent Technologies, Santa Clara, CA, USA). The analyzer performs automatic measurements of the oxygen consumption rate (OCR) and the extracellular acidification rate (ECAR) in real time, being the latter a proxy for lactate formation. Fresh purified GCs were seeded in XF24-well microplates (Agilent Technologies) at a concentration of 3.5×10^5^ cells/well and 3×10^5^ cells/well for CCs. Cells were maintained for 24 hours at 37°C and 5% CO_2_. One hour prior to OCR and ECAR measurements, the supernatant was carefully removed, wells washed with 1 mL of assay medium and, finally, 500 μL of assay medium were added. The assay medium consisted of XF-DMEM medium supplemented with 2% of FBS, 5 mM glucose, 5 mM 4-(2-hydroxyethyl)-1-piperazine ethanesulfonic acid (HEPES), 2 mM glutamine, and 6% FF (see results section) pH 7.4. Cells were incubated at 37°C for 1 hour in a CO_2_-free incubator before loading the plate into the analyzer. The buffering power of the medium was 0.036 mpH units/pmol H^+^ which was determined adding known amounts of HCl to the medium and recording the changes using a pH meter [30].

Control experiments were performed with A549 lung adenocarcinoma cells that were obtained from the American Type Culture Collection. Cells were cultured at 37°C in a humidified 5% CO_2_ atmosphere in DMEM (GIBCO) supplemented with 10% heat-inactivated FBS (Gibco), 2 mM glutamine, and 100 U/mL of penicillin/streptomycin (Invitrogen). A549 cells were seeded in XF24-well microplates at a concentration of 3×10^4^ cells/well and experimental medium was XF-DMEM medium with 2% FBS, 5 mM glucose, 2 mM glutamine, 5 mM HEPES pH 7.4.

The experimental protocol to determine the bioenergetics parameters was essentially as described in [31] which has been often termed as “mitochondrial stress test”. Basal OCR and ECAR were determined by performing four measurements before the addition of the inhibitors or activators (Supplemental Figure 2). Subsequently, ATP turnover was assessed from the decrease in the oxygen consumption after the addition of the ATPase inhibitor oligomycin (2 μM). The ECAR value after the inhibition of mitochondrial ATP synthesis was considered as the maximal ECAR. The maximal respiratory capacity and the spare respiratory capacity were established from the increase in the rate of respiration elicited by the uncoupler carbonyl cyanide *p*-(trifluoromethoxy)-phenylhydrazone (FCCP) after two consecutive additions (0.5 and 0.3 μM). Finally, 1 μM rotenone (complex I inhibitor) and 1 μM antimycin A (complex III inhibitor) were added to quantify the non-mitochondrial respiration. ECAR values were corrected estimating the CO_2_ contribution to the ECAR signal from the correlation between the decreases in the OCR and the ECAR upon the addition of 30 mM 2-deoxyglucose (2-DOG). In order to normalize OCR and ECAR data to take into account differences in cell content, once the experiments finished, wells were washed with PBS and the protein concentration in each well was determined by bicinchoninic acid assay using bovine serum albumin as standard. The protein concentration of CCs could not be determined since the PBS wash led to a partial detachment of the cells and, thus, measurements were unreliable.

### Adenine nucleotides measurement

AMP, ADP and ATP levels were determined by reverse-phase high performance liquid chromatography (HPLC) essentially as described in [32]. A minimum of 4×10^6^ purified GCs were suspended in 10 mL of the same assay medium described in the previous section and incubated at 37°C for 1 hour. Next, they were centrifuged at 8,000 g for 30 sec, homogenized in 660 mM HClO_4_ plus 10 mM theophylline and kept on ice. The homogenates were centrifuged once more at 16,000 g for 15 min at 4°C. The supernatants were neutralized with 2.8 M K_3_PO_4_ until a pH between 6 and 7 was reached and stored at −80°C. Having completed the collection, samples were thawed and centrifuged at 16,000 g for 15 min at 4°C. The supernatants were passed through a 0.45 μm filter and analyzed with a Shimadzu Prominence chromatograph (Canby, Oregon, USA) using a C18 column (Mediterranea SEA18, Teknokroma, Spain). Peaks were identified according to the retention times of standard adenine nucleotides. Chromatograms were analyzed using PeakFit software (version 4.12, Systat Softwate Inc., London, UK). Peak assignment was confirmed using samples treated with 2 μM oligomycin plus 30 mM 2-DOG in which the AMP and ADP peaks markedly increase. The cellular energy charge was calculated using the equation (ATP+ADP/2)/(ATP+ADP+AMP) [33].

### Ethical approval

The Ethics Committee of the University Hospital Puerta de Hierro (Majadahonda, Spain) approved the final version of the study protocol (Identification Code 1512-MAD-064-JG) which also complied with Spanish and European legislation on assisted reproductive technologies. All the included participants signed an informed consent.

### Statistical analysis

Data analysis and graphing were performed using SigmaPlot v14 (Systat Softwate Inc., London, UK) and SPSS v24 (SPSS Inc., Chicago, IL, USA). The continuous variables were reported as mean ± standard deviation (SD) or standard error of the mean (SEM) and compared using the *t*-test or ANOVA, accordingly. The parameters expressed as medians with ranges presented a non-normal distribution and were compared by the Mann-Whitney U test. As appropriate, the categorical variables were analyzed by either the chi-square test or Fisher’s exact test and described as percentages. Statistical significance was set at a two-tailed p-value of < 0.05.

## Results

### Characterization of the bioenergetic profile of human granulosa and cumulus cells

In order to characterize the bioenergetic properties of both mural GCs and CCs, a cohort of oocyte donors aged less than 35 years old was used. The characterization was performed using the XF24 Extracellular Flux Analyzer. Experiments using this technology ought to be designed in culture media supplemented with a set of substrates in order to understand the specific metabolic requirements of the cells. The more common supplements are glucose, pyruvate or glutamine. Depending on the experimental setup, cells will display differences in their bioenergetic profile and associated parameters such as the respiratory capacity or the balance between aerobic glycolysis and oxidative phosphorylation (OXPHOS). The standard conditions commonly used to grow both GCs and CCs include high glucose M-199 media with 10% FBS [34–36]. However, experiments with the XF24 Analyzer were modified to accommodate standard assay conditions including the use of the Seahorse XF DMEM medium buffered with 5 mM HEPES and containing 2% FBS and 2 mM glutamine. Additionally, 5 mM glucose was used to approximate normoglycemic levels.

Initial experiments on the XF24 revealed an unusual profile in which the addition of the uncoupler FCCP caused a modest and transient stimulation of respiration that was followed by a decline in the respiratory rate (Figure 1A). The presence of glutamine and/or pyruvate did not alter the profile significantly (data not shown). Inside the pre ovulatory follicles, cells are immersed in a complex medium, the follicular fluid, containing a variety of metabolites, hormones and other protein factors that are essential for oocyte developmental competence [37–39]. Therefore, we tested if the addition of FF in the assay medium would improve the respiratory capacity and/or the bioenergetic properties of GCs and CCs. Interestingly, the addition of low concentrations of FF (2-6% v/v) markedly increased the response to FCCP in the two cell types (Figure 1B and Supplemental Figure 3). Higher FF concentrations (10%) compromised the adherence of the cells to the plates. Therefore, the standard medium for the subsequent bioenergetic studies always contained the Seahorse XF DMEM medium supplemented with 6% FF, 5 mM glucose, 5 mM HEPES, 2 mM glutamine and 2% FBS. Under these conditions, the stimulation of respiration by FCCP was 107 % ± 6.1 for GCs and 146 % ± 5.9 for CCs. The main results of the characterization of the GCs are summarized in Figure 3. The rate of glycolysis, as determined from the extracellular acidification rate, reveals the high glycolytic activity of the two cell types. Remarkably, ECAR could not be further stimulated when mitochondrial ATP production was inhibited by oligomycin (Figure 2). This was observed in both cell types and it was not influenced by the presence of FF. Their glycolytic profile is also reflected in the OCR/ECAR ratio that is 1.00 ± 0.036 pmol O_2_/mpH unit for GCs and 2.25 ± 0.12 pmol O_2_ /mpH unit for CCs in the presence of 6% FF. Although there is a statistically significant difference between the two types of cells (*p*-value < 0.001), both values correspond to highly glycolytic cells [40].

**Figure 1.**
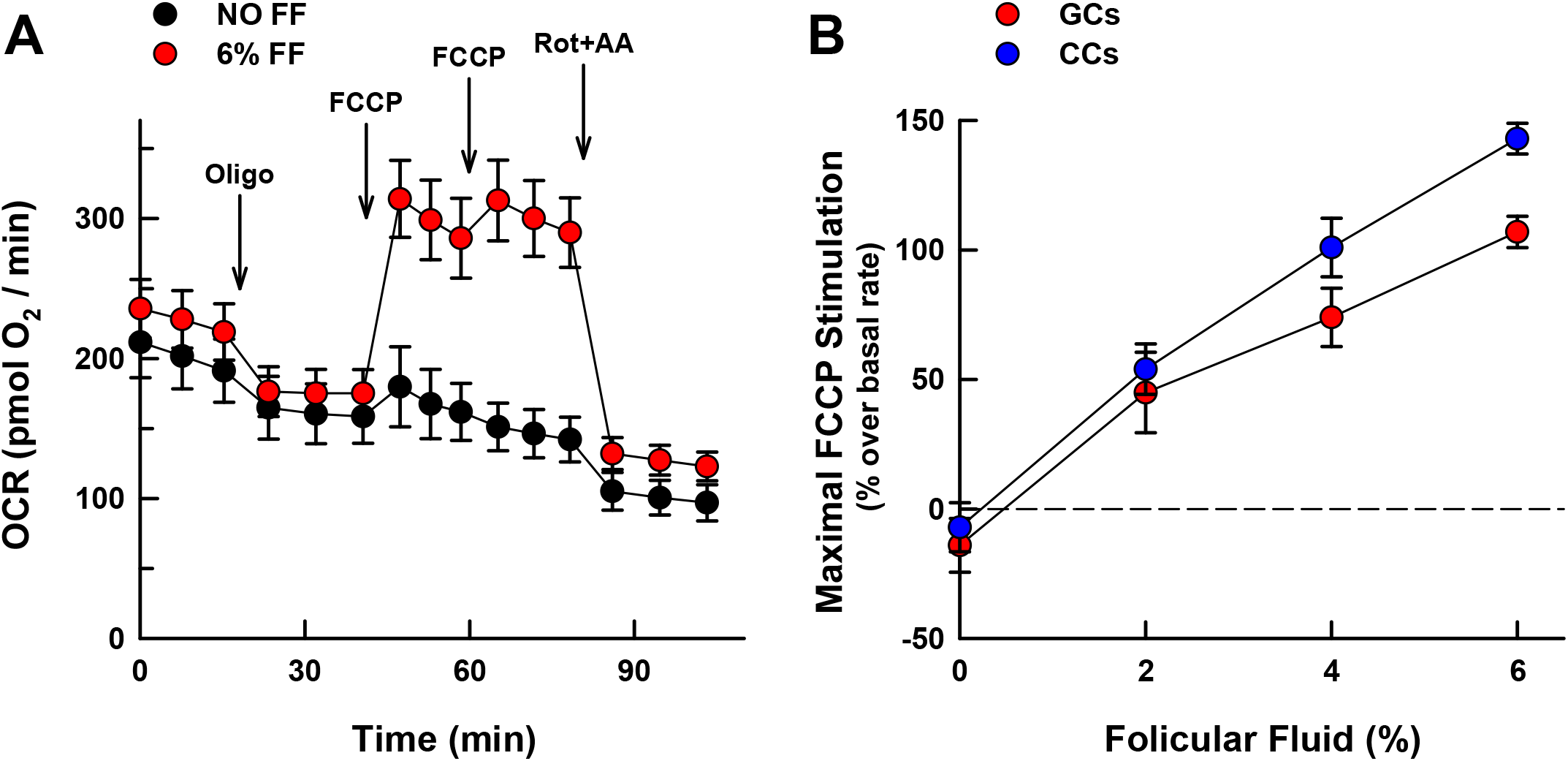
Influence of follicular fluid (FF) on the bioenergetics of granulosa (GCs) and cumulus cells (CCs). (A) Oxygen consumption rates (OCR) of granulosa cells in the absence (black circles) or in the presence (red circles) of 6% FF in the experimental medium. Additions are indicated with the arrows: “Oligo”, 1 μM oligomycin; “FCCP”, two consecutive additions of 0.5 and 0.3 μM; “Rot+AA”, 1 μM rotenone plus 1 μM antimycin A. See Supplemental Figure 2 for further details. (B) Dependence of the maximum respiratory capacity on the presence of FF in GCs (red circles) and CCs (blue circles). The results are presented as mean ± SEM of 6 (GCs + no FF), 4 (GCs + 2% FF), 4 (GCs + 4% FF), 11 (GCs + 6% FF), 4 (CCs + no FF), 4 (CCs + 2% FF), 6 (CCs + 4% FF), and 3 (CCs + 6% FF) independent experiments.

**Figure 2.**
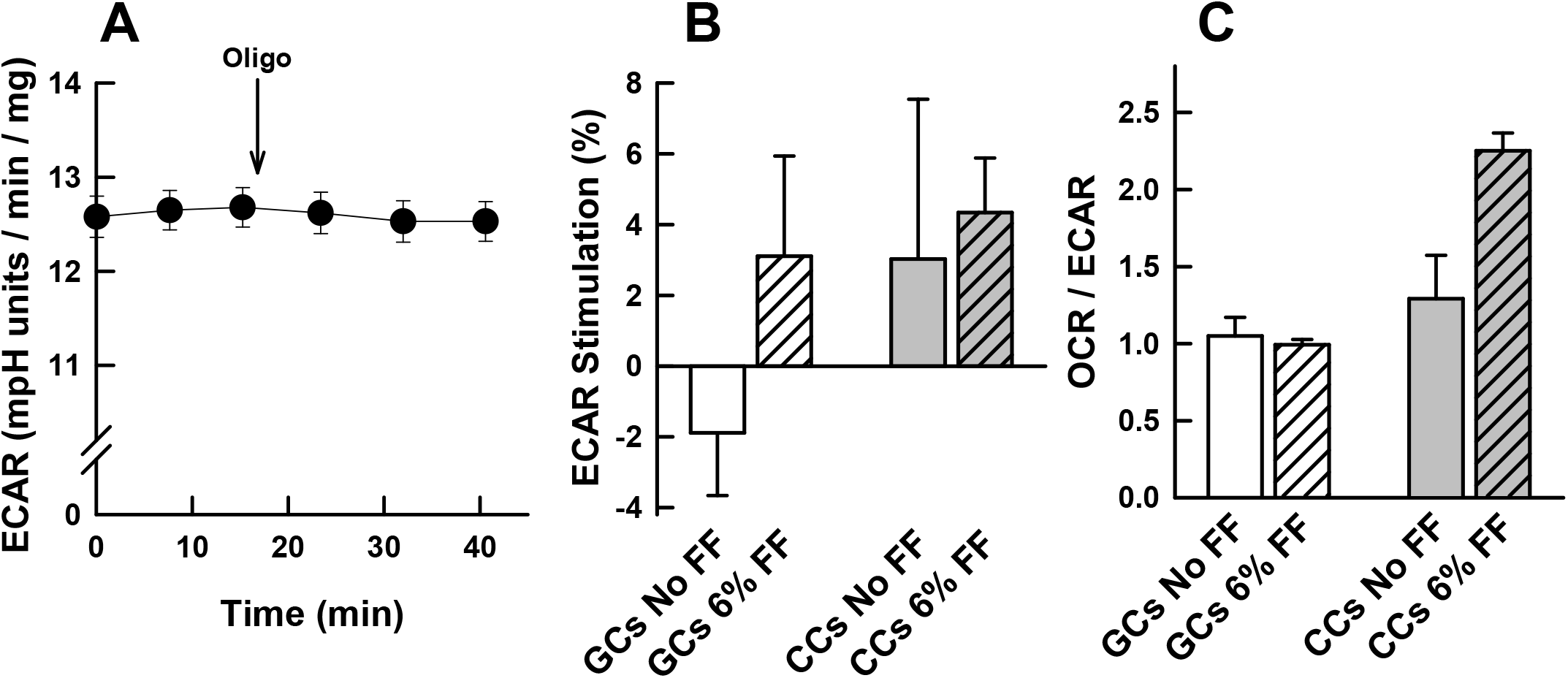
The rate of glycolysis (ECAR) in granulosa (GCs) and cumulus cells (CCs). (A) Effect of oligmycin on the rate of lactate formation (ECAR) of GCs in the presence of 6% follicular fluid (FF). Data points represent the mean ± SEM of 6 independent experiments. (B) Change in ECAR in response to the addition of oligomycin in GCs and CCs both in the presence and absence of FF. Data are expressed as the percent change with respect to basal values. (C) Bioenergetic profile of GCs and CCs expressed as the OCR/ECAR ratio in the presence and absence of FF. Bars represent the mean ± SEM of 11 (GCs + 6% FF), 6 (GCs no FF), 3 (CCs + 6% FF), and 4 (CCs no FF) independent experiments.

**Figure 3.**
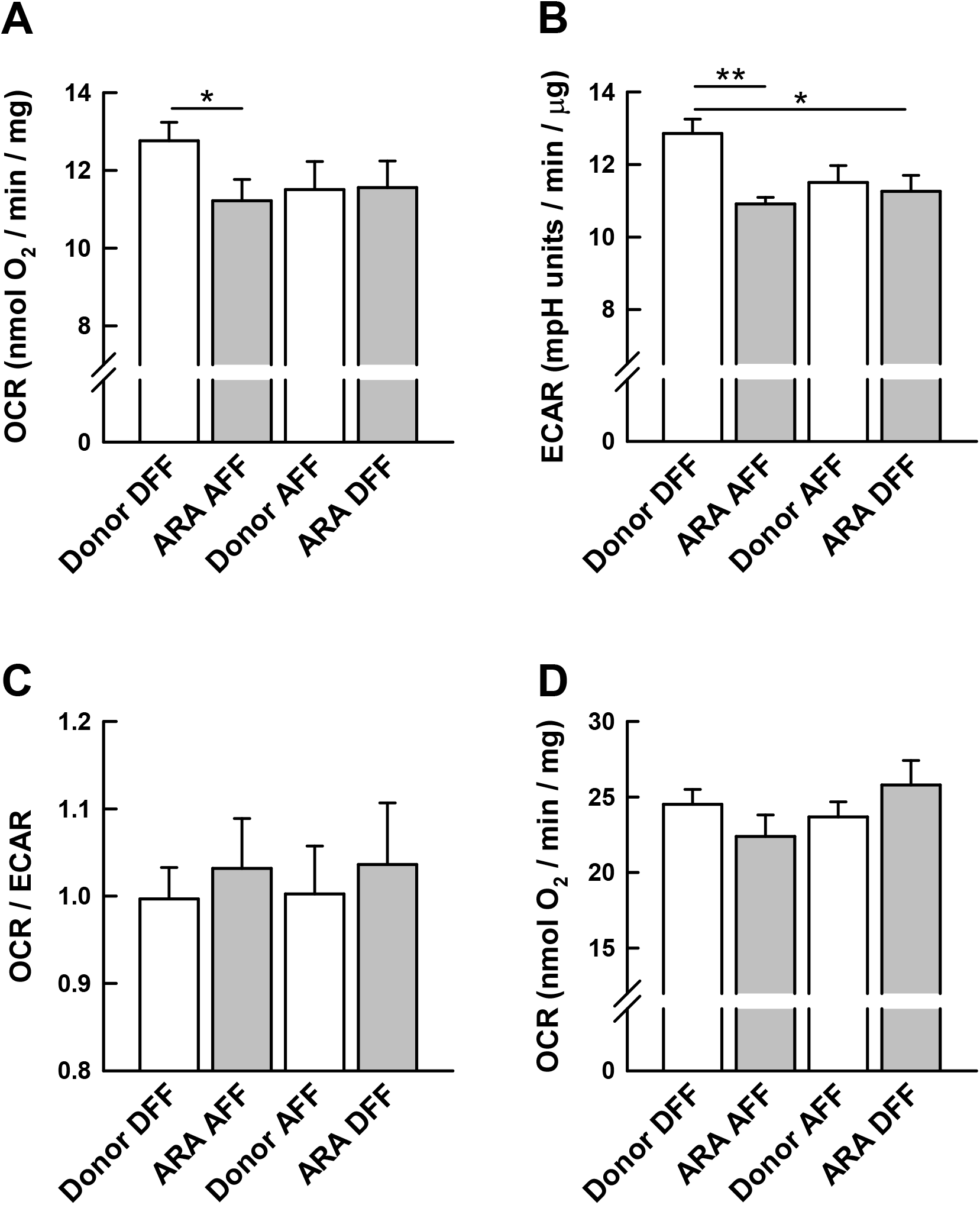
Bioenergetic parameters of mural granulosa cells (GCs) from young donors (empty bars) and advanced reproductive age (ARA) women (grey bars) and influence of the FF origin: DFF, FF from donors; AFF, FF from ARA group. (A) Basal rate of mitochondrial respiration. (B) Basal rate of glycolysis as determined from the rate of lactate formation (ECAR) value. (C) Bioenergetic profile expressed as the OCR/ECAR ratio. (D) Maximum respiratory capacity. Bars represent the mean ± SEM of 11 (Donor DFF and ARA DFF) or 9 (Donor AFF, ARA AFF) independent experiments. **P<0.01, *P<0.05 between the indicated groups (ANOVA test).

The bioenergetic profile of mural GCs and CCs presented an intriguingly high component of non-mitochondrial respiration which should be due to oxygen consuming processes like those catalyzed by cyclooxygenases, lipoxygenases or NADPH oxidases (NOX). In most cellular systems the non-mitochondrial OCR generally accounts for less than 10% of the basal OCR [31] (the bioenergetic profile of A549 cells is shown in Supplemental Figure 4A for a comparison). There are important exceptions like the activity of non-mitochondrial NOX in macrophages that can contribute significantly to cellular oxygen uptake. Under our experimental conditions, the non-mitochondrial respiration in GCs accounted for 56% ± 1.1 of the basal OCR. The NOX inhibitors apocynin and VAS2870 had no effect on the non-mitochondrial respiration of GCs thus ruling out the involvement of NOX (Supplemental Figure 4B).

### Effects of senescence on the energy metabolism of granulosa cells

Since female fertility and IVF outcomes dramatically decline with age, a study was designed to explore potential changes in the energy metabolism of GCs from a group of patients of advanced reproductive age (ARA). The study was restricted to mural GCs because the yield of cells from each patient was significantly higher and, as previously stated, the attachment of the CCs to the XF24 plate was weaker. Patients’ characteristics, cycle parameters and ovarian response to COS are shown in Table 1. The mean age of the ARA group was 40.8 ± 2 while in the control group it was 24.1 ± 3.9. Therefore, the difference between the two experimental groups allowed exploring the effect of age-related infertility on the bioenergetics of human GCs. The mean level of anti-Müllerian hormone (AMH) in the ARA group was 1.27 ± 0.95 ng/ml. Egg donors AMH levels are not routinely quantified in our Institution, however the antral follicle count (AFC) was remarkably higher in this group (19 ± 4.4 *vs.* 9 ± 4.2; *p*-value < 0.001). The ARA group required higher gonadotropin doses [2000 IU (775-4800) *vs.* 2250 IU (1350-4275); *p*-value < 0.001]. As expected, the number of oocytes retrieved was significantly higher in the egg donors’ group [21 ± 9.2 *vs*. 9 ± 5.7; *p*-value < 0.001], as was the number of mature oocytes [14 (5-41) *vs*. 6 (1-23); *p*-value < 0.001)].

Knowing the influence of the FF on GC bioenergetics, the assay media for the control and ARA patients were supplemented with FF isolated from individuals belonging to their respective groups. It could be expected that the observed bioenergetic properties could be conditioned by the cells, the composition of the FF or a combination of the two factors. Figure 3 shows a summary of the bioenergetic properties of GCs from the control donors and the ARA group. It is apparent that the basal rate of respiration and the ECAR are significantly reduced in the ARA group thus indicating a global decrease in the energy metabolism as woman age. As expected, the parallel decrease in the two parameters does not modify the bioenergetic profile as estimated by the OCR/ECAR ratio (Figure 3C). Notably, the decrease in the basal respiration in the ARA group is not reflected in the maximal respiratory capacity of the cells, which remains unchanged (Figure 3D).

The influence of the origin of the FF used to supplement the media was investigated performing experiments in which the donor FF was used to supplement the media for the ARA group and *viceversa*. Figure 3 reveals that the highest rates of respiration and glycolysis were observed in the donor group with medium supplemented with the donor FF. The rest of combinations led to a decrease in the energy metabolism although under no condition the maximal respiratory capacity was affected.

### Adenine nucleotides levels in human GCs

To investigate if the decreased metabolism is due to senescence and causes a reduction in the cellular energy charge, the adenine nucleotide levels (ATP, ADP and AMP) of the GCs in the two experimental groups were determined. The incubation media was the same than in the XF24 experiments, supplementing the XF-DMEM medium with 6% FF of the respective experimental group. Figure 4 summarizes the changes in the adenine nucleotide pool and reveals that GCs from the ARA group exhibited a marked decrease in the cellular energy charge, which is also reflected in the ATP/ADP and ATP/AMP ratios. Since experiments on the XF24 analyzer had shown that inhibition of mitochondrial ATP synthesis did not cause a compensatory increase in glycolysis (ECAR), the effect of oligomycin on the adenine nucleotide pool was studied. Results in Figure 4 (hatched bars) show that this inhibitor markedly decreases ATP levels in both groups confirming that, since glycolysis cannot be further stimulated, it cannot compensate for the loss of the mitochondrial ATP synthesis. As expected, the combination of oligomycin and 2-DOG totally collapsed the energy levels.

**Figure 4.**
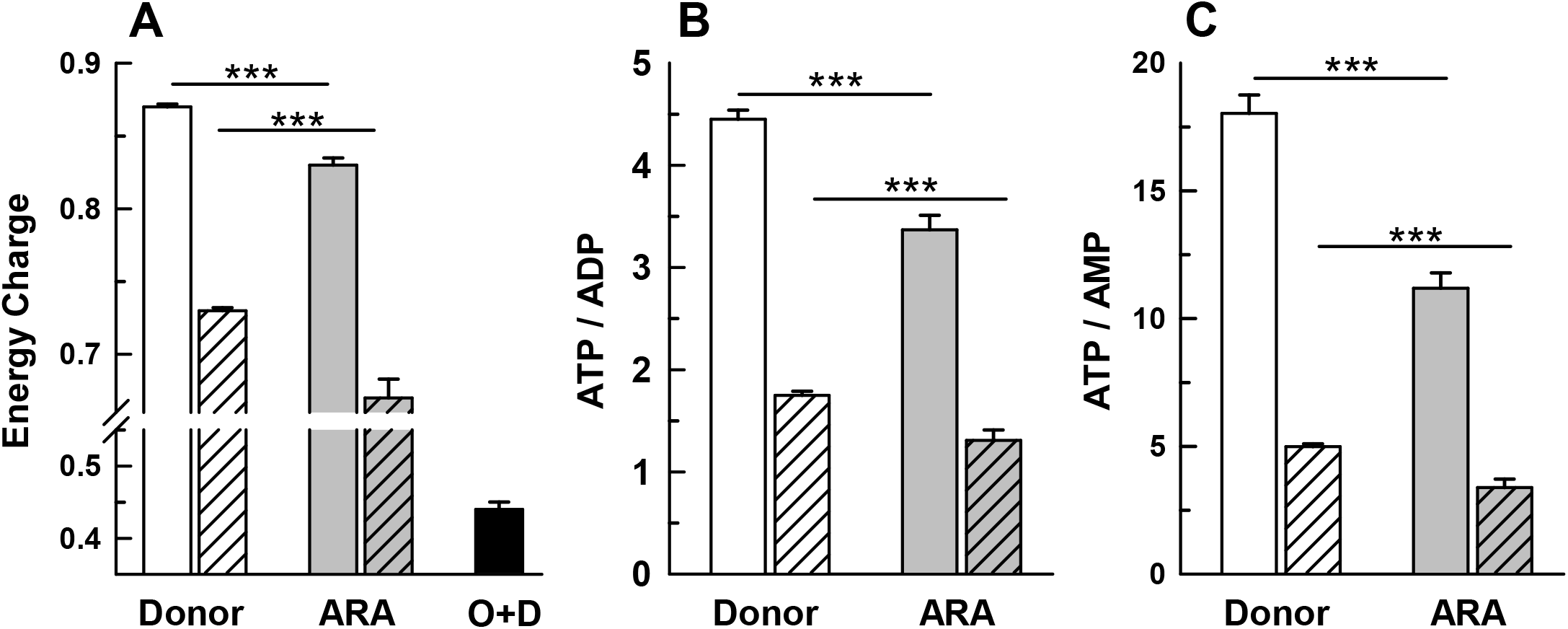
Effect of reproductive senescence on the adenine nucleotide pool in mural granulosa cells (GCs) in the presence of 6% follicular fluid (FF). Empty bars, control donors. Grey bars, ARA group. Hacthed bars in each group are the values obtained in the presence of oligmycin. Black bar is the value in the presence of oligomycin and 2-deoxyglucose. (A) Energy charge calculated using the equation (ATP+ADP/2)/(ATP+ADP+AMP). (B) ATP/ADP ratio. (C) ATP/AMP ratio. Bars represent the mean ± SEM of 14 (Donor), 9 (ARA), 7 (Donor + oligomycin), 5 (ARA + oligomycin), 7 (Donor + oligomycin + deoxyglucose) independent determinations. ***P<0.001 between the indicated groups (ANOVA test).

## Discussion

The analysis of the bioenergetic properties of human mural GCs and CCs presented have allowed the progress in the understanding of the metabolism of these two follicular cell types that are known to influence the maturation of the oocyte and, ultimately, IVF outcomes [41–42]. These cells are believed to be potential biomarkers of oocyte quality [39,43], and it has been proposed that alterations in their energy metabolism could lead to infertility [27,41,42,44]. The results evidence (i) the high glycolytic profile of these two cell types and (ii) that the respiratory capacity relies on the presence of follicular fluid in the experimental media (Figure 1 and Supplemental Figure 3). When evaluating oxygen consumption rates, we observed a strikingly high non-mitochondrial OCR that does not appear related to NOX activity and deserves further investigation. Among the various oxygen-consuming enzymes, the involvement of cyclooxygenase-2 in prostaglandin biosynthesis in GCs should be considered since its major product, the prostaglandin E2 (PGE2), is an important mediator of ovulation and it is presumably increased during gonadotropins stimulation [45,46].

The follicular fluid is a complex medium produced by GCs and thecal cells, and also enriched with biomolecules from plasma, that influences oocyte competence and quality [39,47,48]. The main components are steroid hormones, metabolites (amino acids, glucose, fatty acids, etc.), polysaccharides, proteins, and growth factors among others [37–39]. The presence of enzymatic and non-enzymatic antioxidants is essential to maintain homeostatic levels of reactive oxygen species that are potentially harmful but also necessary for oocyte growth [48,49]. Although our data has revealed that the FF is a fundamental element to attain the full respiratory capacity of mural GCs and CCs, it was beyond the scope of the current work to elucidate which are the critical components.

The respiratory capacity of the cells, under our experimental conditions, was determined by adding an artificial uncoupling agent (FCCP) and, therefore, its physiological relevance should be interpreted with caution. However, if this bioenergetic parameter were to reflect the OXPHOS capacity of the oocyte and/or the supporting cells, then FF components could significantly affect the outcome of the energy demanding processes that take place upon fertilization.

The other pathway that provides energy to the cell is glycolysis. Our data show that, under our experimental conditions, the rate of glycolysis appears to be at its maximum since the inhibition of mitochondrial ATP synthesis did not result in a compensatory increase in glycolysis (Figure 2) and, consequently, inhibition of mitochondrial ATP synthesis led to a collapse in the cellular energy charge (Figure 4). These findings emphasize the importance of the two pathways to maintain an adequate ATP supply and sustain development. The decrease in both OXPHOS and glycolysis could be the result of a decreased energy demand of the cell due to reproductive senescence. If this were to be the case, the cellular energy charge should remain unchanged. However, our data clearly shows that aging impairs cellular bioenergetics (Figure 3) and as a result there is a significant decrease in the ATP levels (Figure 4). It is puzzling that despite the lower energy charge observed in the ARA group, OXPHOS is not increased to attempt to restore the optimal ATP levels despite the existence of spare respiratory capacity. In this context, it must be pointed out that the maximal respiratory capacity remains unchanged in the two experimental groups. This is important since it reveals that the mitochondrial mass (total respiratory capacity) does not appear to be affected by aging. A recent study, that used flow cytometry to evaluate mitochondrial mass and respiratory capacity of CCs, failed to detect significant differences between patients with less than 35 and more than 35 years of age [43]. Therefore, the molecular basis for the diminished energy metabolism cannot be envisaged at present. However, there are reports showing, for example, that in human GCs deficiencies in complex V [42] or variations in sirtuin expression [50] could be behind mitochondrial dysfunction in ovarian aging.

Compelling evidence suggests that the composition of the human FF plays a role in IVF outcomes [38,51,52], however, the defects in energy metabolism during reproductive senescence are only partially explained by its composition. Thus, basal rates of respiration and glycolysis in GCs from the control group were significantly reduced in the presence of FF from older women but FF from the young donors fail to improve the bioenergetic parameters in the ARA group (Figure 3). Recently, a porcine in-vitro maturation model showed that supplementing the maturation medium with FF not only increased the mitochondrial DNA copy number and the ATP content in both oocytes and cumulus cells, but also the survival of cumulus cells and the blastulation rate [53].

A word of caution should be included regarding the physiological significance of these observations since it cannot be ascertained if they can be extrapolated to GCs or CCs from growing and non-preovulatory follicles. There is a limitation in the experimental approach, derived from the established IVF protocols, which implies that luteinized cells from preovulatory follicles can only be collected following oocyte retrieval. Therefore, in humans, these issues will only be approached when *in vitro* maturation protocols start to be widely applied in fertility care units.

In conclusion, many reports have pointed out to mitochondria as playing a critical role in oocyte quality and IVF outcome [8–11]. However, we have shown that both OXPHOS and glycolysis are decreased in mural GCs during reproductive aging in women and these two concerted events lead to a decrease in ATP levels. Moreover, mural GCs retain their full respiratory capacity although unknown factors prevent the expected increase in mitochondrial respiration that should restore the cellular energy charge. It must be pointed out that all samples were obtained from women that have gone through stringent exclusion criteria to ensure that there were no underlying pathologies that could influence fertility. Therefore, it seems reasonable to assume that the observed deficiency in the energy metabolism is related to intrinsic functional properties due to reproductive aging. Such detrimental effects of age on GCs bioenergetics are likely to influence overall IVF performance. A new window of opportunity for diagnostic and therapeutic tools may arise from studies focusing on the bioenergetics of granulosa cells, oocytes and embryos.

## Conflict of interest

The authors declare no conflicts of interest.

## Author contributions

JAGV and ER designed the study. GNC performed sample collection, experiments and statistical analyses. GNC and ER designed the experiments, analysed the data and wrote the manuscript. All the authors approved the final version of the manuscript.

## Abbreviations

2-DOG: 2-deoxyglucose
ARA: advanced reproductive age
CC: Cumulus cell
ECAR: Extracellular acidification rate
FBS: heat-inactivated fetal bovine serum
FCCP: carbonyl cyanide p-(trifluoromethoxy)-phenylhydrazone
FF: Follicular fluid
GC: granulosa cell
HEPES: 4-(2-hydroxyethyl)-1-piperazine ethanesulfonic acid
IVF: *in vitro* fertilization
NOX: NADPH oxidase
OCR: Oxygen consumption rate
OS: Ovarian stimulation
OXPHOS: oxidative phosphorylation

## Notes

**Grant support:** This work was supported by IVI-Madrid. GNC received grants from the Capes Foundation, Brazil/PDSE/Process number 88881.132905/2016-01.

### Competing Interest Statement

The authors have declared no competing interest.

