## Supplemental material for "REPRODUCTIVE SENESCENCE IMPAIRS THE ENERGY METABOLISM OF HUMAN GRANULOSA CELLS"

### Supplementary material

**Supplemental Figure 1.** Graphic summary of the experimental design of the study.

**Supplemental Figure 2.** Scheme of the experimental protocol for the bioenergetic characterization of the cells termed "mitochondrial stress test". The rate of oxygen consumption (blue trace) is monitored while a sequence of OXPHOS effectors are addded: "Oligo", 1 μM oligomycin to inhibit mitochondrial ATP synthesis; "FCCP", two consecutive additions of the uncoupler FCCP (0.5 and 0.3 μM) are made to ensure that the maximum rate of respiration is reached; "Rot+AA", simultaneous addition of 1 μM rotenone and 1 μM antimycin A to inhibit mitochondrial respiration. The following bioenergetic parameters are calculated for the rates of oxygen consumption as indicated in the scheme: 1, Basal rate of mitochondrial respiration; 2, Non-mitochondrial oxygen consumption; 3, Maximum rate of respiration also termed as "respiratory capacity"; 4, Spare respiratory capacity; 5, ATP turnover (31).

**Supplemental Figure 3**. Effect of the follicular fluid (FF) on the bioenergetics of (A) mural granulosa cells (GCs) and (B) cumulus cells (CCs). Mitochondrial stress tests performed in the presence of increasing concentrations of FF. Respiration is normalized to 100% in the OCR measurement just before oligomycin. Additions: "Oligo", 1 μM oligomycin; "FCCP", two consecutive additions of 0.5 and 0.3 μM; "Rot+AA", 1 μM rotenone plus 1 μM antimycin A. Data points represent the mean ± SEM of 6 (GCs + no FF), 4 (GCs + 2% FF), 4 (GCs + 4% FF), 11 (GCs + 6% FF), 4 (CCs + no FF), 4 (CCs + 2% FF), 6 (CCs + 4% FF), and 3 (CCs + 6% FF) independent experiments.

**Supplemental Figure 4**. Analysis of the non-mitochondrial respiration in mural granulosa cells (GCs). (A) Comparison between the profile of the "mitochondrial stress tests" performed with GCs (red circles) and the human adenocarcinoma cell line A549 (black circles). For this comparison two assays with similar basal respiration were chosen to illustrate the difference in the non-mitochondrial respiration. (B) Lack of effect of the NADPH oxidases (NOX) inhibitors VAS2870 10-20 μM and apocynin 500 μM on the non-mitochondrial respiration. The inhbitors were added 60 min before the experiment.

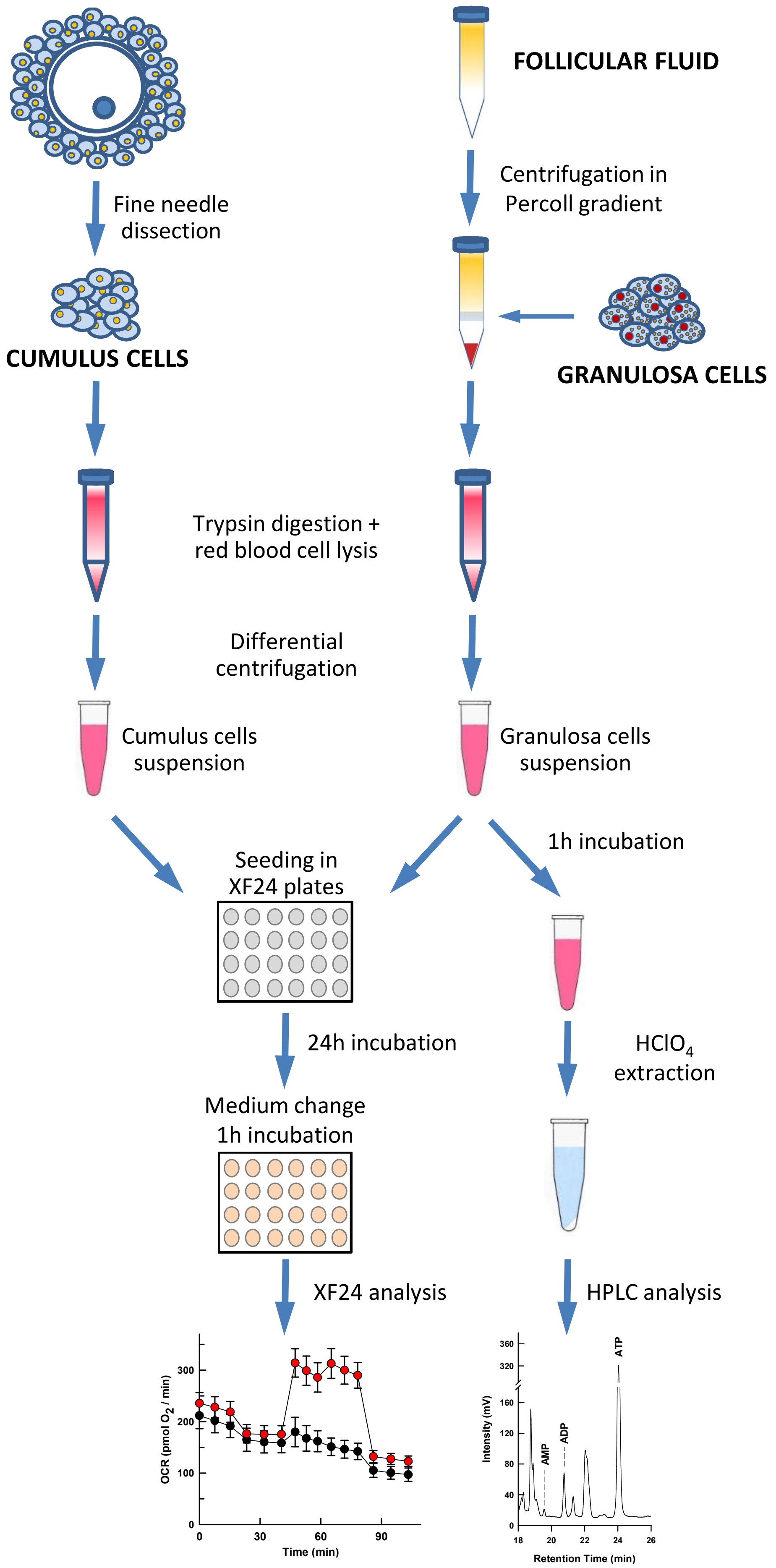


Cecchino *et al.* Supplemental Figure 1

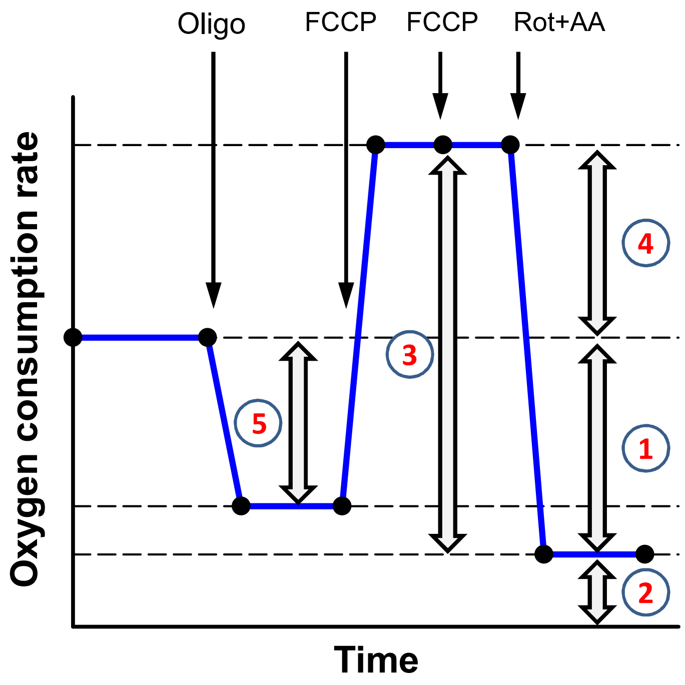


Cecchino *et al.* Supplemental Figure 2

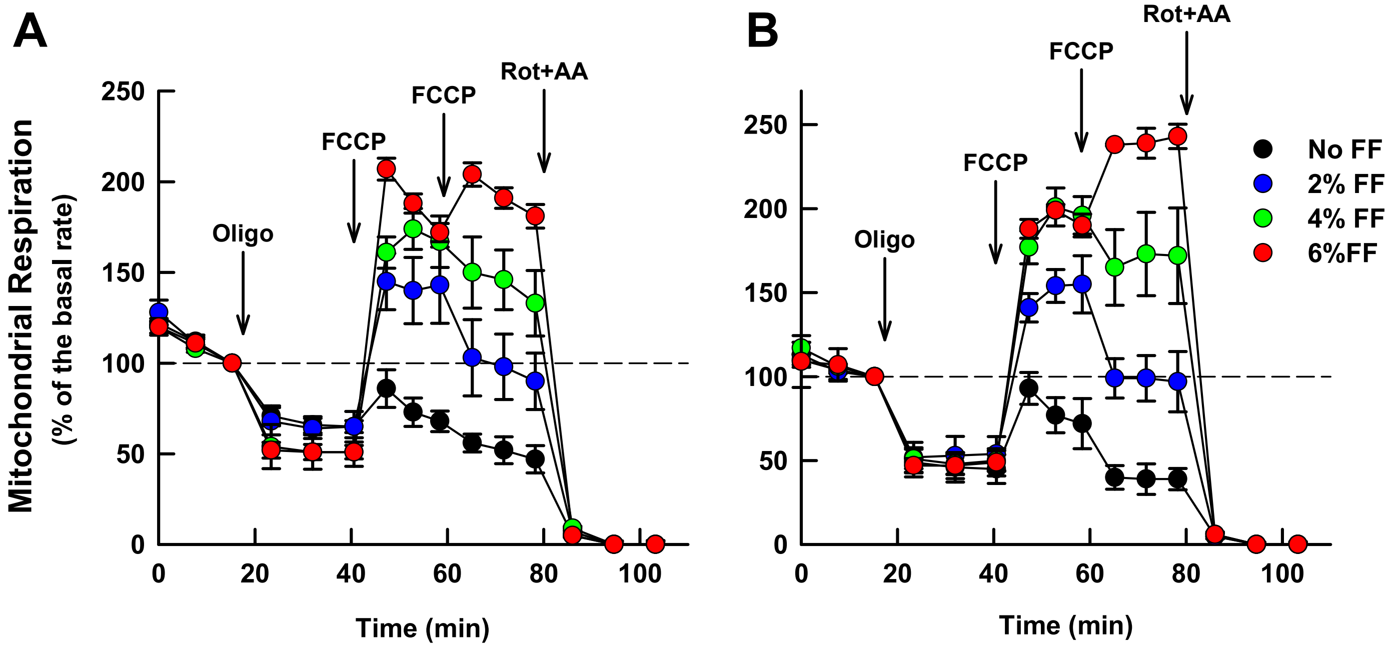


Cecchino *et al.* Supplemental Figure 3

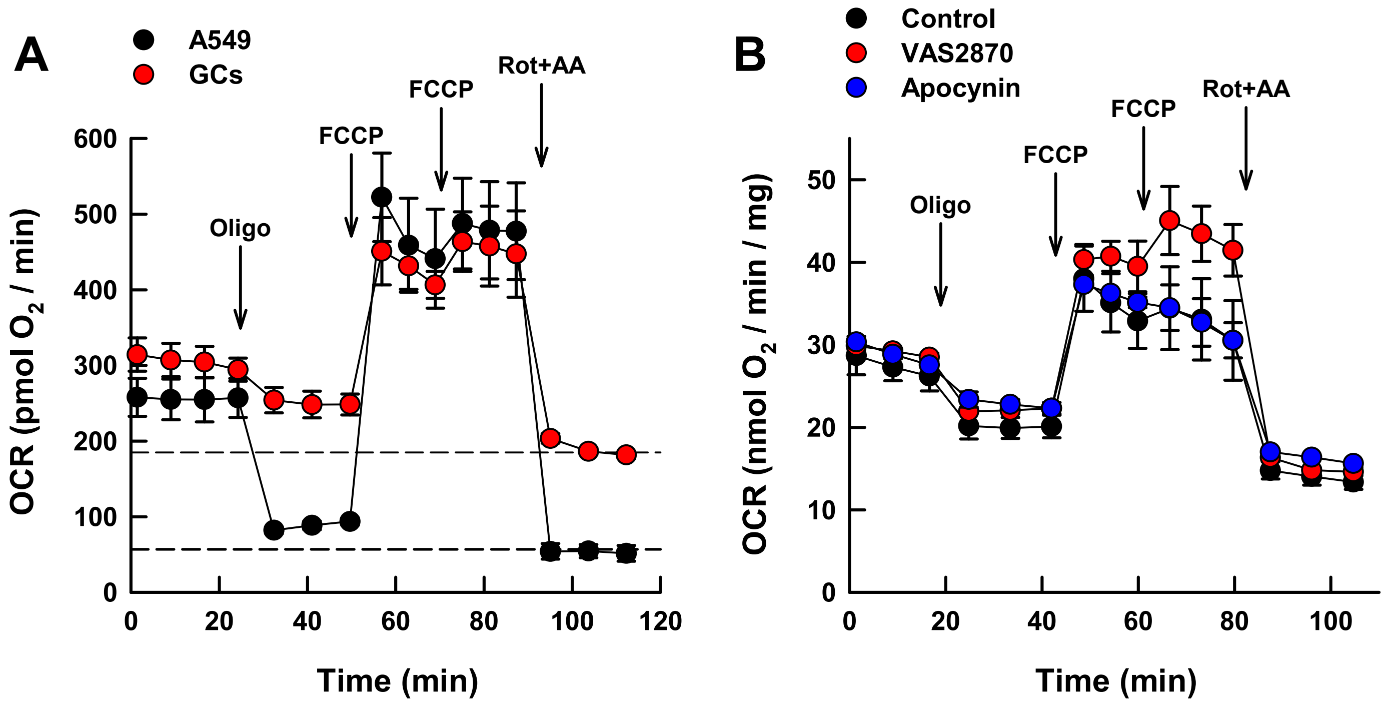


Cecchino *et al.* Supplemental Figure 4
